# Gradual centriole maturation associates with the mitotic surveillance pathway in mouse development

**DOI:** 10.1101/2020.07.24.219402

**Authors:** Cally Xiao, Marta Grzonka, Charlotte Gerards, Miriam Mack, Rebecca Figge, Hisham Bazzi

## Abstract

Centrosomes, composed of two centrioles and pericentriolar material, organize mitotic spindles during cell division and template cilia during interphase. The first few divisions during mouse development occur without centrioles, which form around embryonic day (E) 3. However, disruption of centriole biogenesis in *Sas-4* null mice leads to embryonic arrest around E9. Centriole loss in *Sas-4*^−/−^ embryos causes prolonged mitosis and p53-dependent cell death. Studies *in vitro* discovered a similar USP28-, 53BP1-, and p53-dependent mitotic surveillance pathway that leads to cell cycle arrest. In this study, we show that an analogous pathway is conserved *in vivo* where 53BP1 and USP28 are upstream of p53 in *Sas-4*^−/−^ embryos. The data indicates that the pathway is established around E7 of development, four days after the centrioles appear. Our data suggest that the newly formed centrioles gradually mature to participate in mitosis and cilia formation around the beginning of gastrulation, coinciding with the activation of mitotic surveillance pathway upon centriole loss.

## Introduction

Centrosomes are major microtubule organizing centers (MTOCs) of animal cells and are composed of two centrioles, one mature mother centriole with distal and sub-distal appendages and one daughter centriole, surrounded by a proteinaceous pericentriolar material (PCM) (Conduit *et al*, 2015). During mitosis, centrosomes help assemble the mitotic spindle, and during interphase, the mother centriole forms the basal body template for cilia (Bornens, 2012). In proliferating cells, centrioles can form *de novo* without pre-existing centrioles or use the scaffold of existing centrioles to duplicate once per cell cycle in late G1 and S phases (Loncarek & Khodjakov, 2009). The centriole formation pathway has been defined in cell culture and in different organisms and relies on a set of core proteins that include spindle assembly defective protein 4 (SAS-4, also called CENPJ or CPAP) (Kirkham *et al*, 2003; Kleylein-Sohn *et al*, 2007; Leidel & Gonczy, 2003; Tang *et al*, 2009). The newly formed centrioles undergo maturation over two cell cycles to acquire appendages, become MTOCs and template cilia (Kong *et al*, 2014). Cilia formation relies on docking of the mother centriole to the plasma membrane through distal appendage proteins, such as CEP164 (Graser *et al*, 2007; Siller *et al*, 2017), and on intra-flagellar transport proteins, such as IFT88 (Haycraft *et al*, 2007).

During rodent development, and unlike the development of most organisms, the first cell divisions post-fertilization occur without centrioles (Courtois *et al*, 2012; Gueth-Hallonet *et al*, 1993; Howe & FitzHarris, 2013; Manandhar *et al*, 1998; Woolley & Fawcett, 1973). In the mouse embryo, centrioles first form by *de novo* biogenesis starting at the blastocyst stage around embryonic day (E) 3.5 (Courtois & Hiiragi, 2012). Before centriole formation, diffuse γ-tubulin signals, a PCM component and microtubule nucleator, appear at the morula stage around E3, and γ-tubulin signals become more focused as centrioles form; however, the newly formed centrioles do not seem to act as MTOCs in interphase cells (Howe & FitzHarris, 2013). In addition, the first cilia form almost two days post-implantation around E6.5 in cells of the epiblast (Bangs *et al*, 2015).

Mouse embryonic stem cells (mESCs) are a well-established *in vitro* model of embryo development that are derived from the pluripotent inner cell mass of blastocysts at E3.5 but molecularly resemble epiblast cells post-implantation (Nichols & Smith, 2011). To maintain uniform pluripotency, mESCs are cultured with leukemia inhibitory factor (LIF) and two other differentiation inhibitors abbreviated as 2i (Williams *et al*, 1988; Ying *et al*, 2008). In pluripotency, the transcription factor NANOG is highly expressed in mESCs and regulates self-renewal (Rosner *et al*, 1990). In this study, we used mESCs to complement our *in vivo* experiments by studying the growth dynamics of cells without centrioles.

We have previously shown that the genetic removal of SAS-4 in the mouse resulted in the loss of centrioles and cilia (Bazzi & Anderson, 2014a). The *Sas-4*^−/−^ embryos arrested development around E9.5 due to p53-dependent cell death. The increase in p53 in *Sas-4*^−/−^ embryos was not due the secondary loss of cilia, DNA damage or chromosome segregation errors. Also, these phenotypes are not specific to *Sas-4*^−/−^ embryos because mutations in different genes, such as *Cep152*, that cause centriole loss show similar phenotypes (Bazzi & Anderson, 2014a, b). Notably, the fraction of mitotic cells was higher in *Sas-4*^−/−^ embryos at E7.5 and E8.5, indicating a longer mitotic duration of cells without centrioles, which was also confirmed by time-lapse imaging of dividing cells. Because a short nocodazole treatment to prolong mitosis upregulated p53 in cultured wild-type (WT) embryos, the data suggested that the less efficient mitosis without centrioles activated a novel p53-dependent pathway (Bazzi & Anderson, 2014a). In cultured mammalian cell lines *in vitro*, a similar pathway that is activated by the loss of centrioles or prolonging mitosis leads to p53-dependent cell cycle arrest and is called the mitotic surveillance pathway (Lambrus & Holland, 2017; Lambrus *et al*, 2015; Wong *et al*, 2015). Recently, p53-binding protein 1 (53BP1) and ubiquitin specific peptidase 28 (USP28) have been shown to be essential for the conduction of this pathway *in vitro* (Fong *et al*, 2016; Lambrus *et al*, 2016; Meitinger *et al*, 2016). These studies showed that mutations in *53BP1* or *USP28* rescued the growth arrest phenotype observed in cells without centrioles. However, whether a similar 53BP1- and USP28-dependent pathway operates *in vivo* and can cause the p53-dependent cell death phenotype in the mouse are still not known.

In this study, our data showed that the mitotic surveillance pathway is conserved in mice *in vivo* and that 53BP1 and USP28 are essential for its conduction upstream of p53. In order to explain the late onset of the phenotype upon the loss of centrioles, we also asked when during development this pathway is established. The data indicated that the newly formed centrioles around E3 are not fully mature and do not seem to be required for mitosis until around E7 of development, when the pathway is initiated. Our data suggest that once the cells start to depend on centrosomes as MTOCs in mitosis and ciliogenesis, then they sense the loss of centrioles and activate the p53-dependent mitotic surveillance pathway.

## Results and Discussion

### Mutations in *53bp1* or *Usp28* rescue the *Sas-4* mutant phenotype *in vivo*

To test the conservation of the mitotic surveillance pathway and the involvement of 53BP1 and USP28 in its activation *in vivo, 53bp1*^*+/*−^ and *Usp28*^*+/*−^ null mouse alleles were generated using CRISPR/Cas9 gene editing (see Methods and Fig. EV1) and crossed to *Sas-4*^*+/*−^ mice (Bazzi & Anderson, 2014a). Both *Sas-4*^−/−^ *53bp1*^−/−^ and *Sas-4*^−/−^ *Usp28*^−/−^ embryos showed remarkable rescues of the morphology and size compared to the *Sas-4*^−/−^ embryos at E9.5 (Fig. 1A). *Sas-4*^−/−^ *53bp1*^−/−^ and *Sas-4*^−/−^ *Usp28*^−/−^ embryos both underwent body turning, had visible somites and open heads, and were similar to *Sas-4*^−/−^ *p53*^−/−^ mutants (Bazzi & Anderson, 2014a). At the molecular level, *Sas-4*^−/−^ *53bp1*^−/−^ and *Sas-4*^−/−^ *Usp28*^−/−^ embryos showed highly reduced levels of p53 and cleaved-Caspase 3 (Cl-CASP3) compared to *Sas-4*^−/−^ embryos (Fig. 1B, C). The data indicated that mutating *53bp1* or *Usp28* suppressed both p53 stabilization and p53-dependent cell death upon centriole loss *in vivo* and established the conservation of the mitotic surveillance pathway in the mouse.

**Fig. 1.**
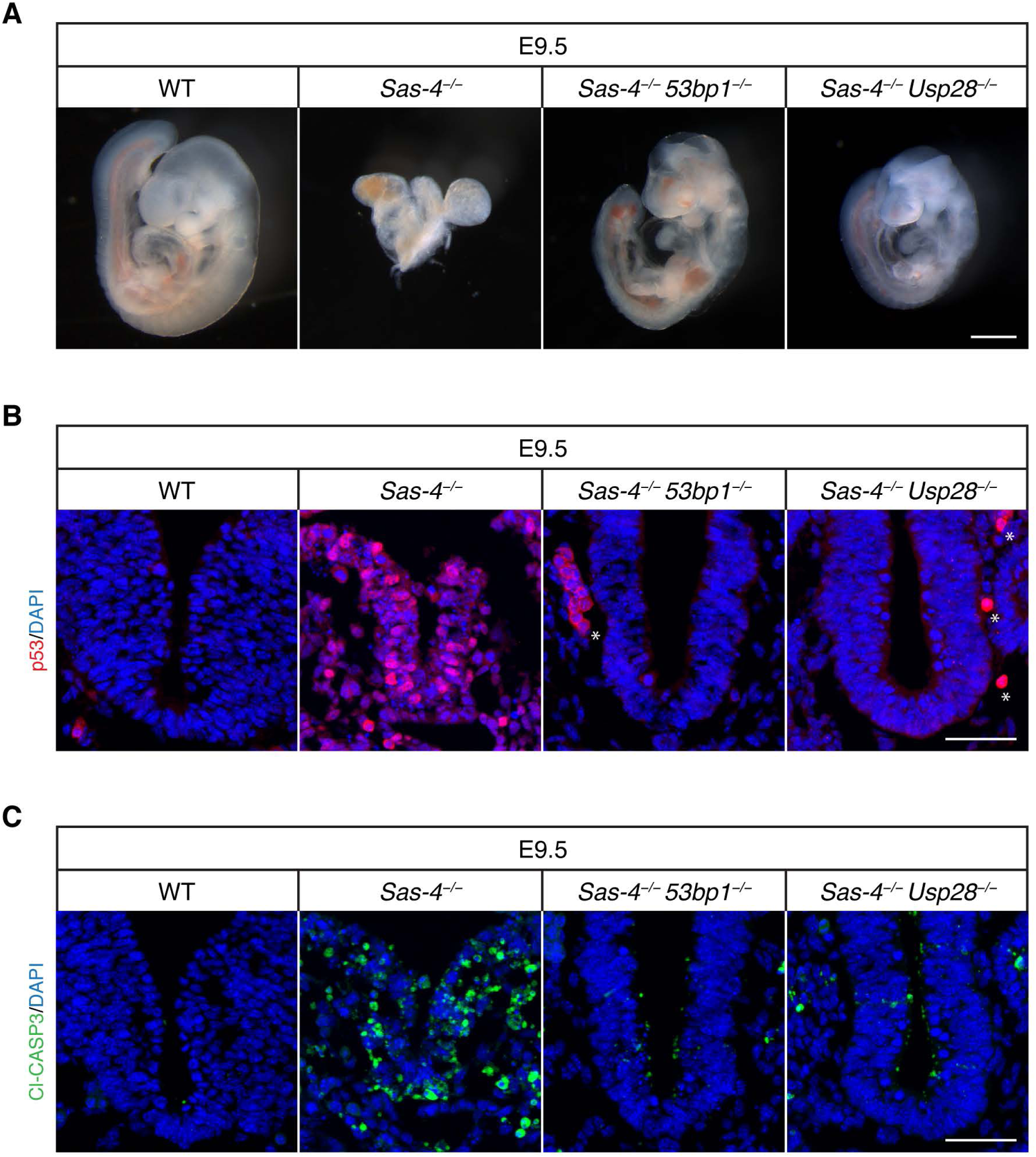
The mitotic surveillance pathway is conserved in the mouse *in vivo*. (A) Gross morphology of WT, *Sas-4*^−/−^, *Sas-4*^−/−^ *53bp1*^−/−^, and *Sas-4*^−/−^ *Usp28*^−/−^ embryos at E9.5. Scale bar = 500 μm. (B, C) Immunostaining for p53 (B) and Cleaved-Caspase3 (Cl-CASP3, C) on transverse sections of WT, *Sas-4*^−/−^, *Sas-4*^−/−^ *53bp1*^−/−^, and *Sas-4*^−/−^ *Usp28*^−/−^ embryos at E9.5. The area shown encompasses the ventral neural tube and surrounding mesenchyme. Asterisks in B denote non-specific staining of blood cells. Scale bar = 50 μm.

### The mitotic surveillance pathway is activated around E7

In order to determine when the mitotic surveillance pathway is activated in *Sas-4*^−/−^ embryos, we used immunostaining and quantified nuclear p53 levels during development. At E7.5, *Sas-4*^−/−^ embryos were smaller than control embryos (WT or *Sas-4*^*+/*−^) with around 1.5-fold higher nuclear p53 in the epiblast (Fig. 2A, B) (Bazzi & Anderson, 2014a). Earlier in development at E6.5, *Sas-4*^−/−^ embryos were morphologically indistinguishable from control embryos, and nuclear p53 was not detectably different (Fig. 2A, B). *Ift88* null (cilia mutant) embryos were used as controls for *Sas-4*^−/−^ centriole mutant embryos, and were similar to WT embryos both morphologically and in terms of p53 nuclear levels at E7.5 and E6.5 (Fig. EV2), confirming our earlier finding that p53 upregulation was due to centriole loss and not the secondary loss of cilia (Bazzi & Anderson, 2014a). The data suggested that the increased level of nuclear p53 in *Sas-4*^−/−^ embryos starts around E7 of development and is independent of cilia loss *per se*.

**Fig. 2.**
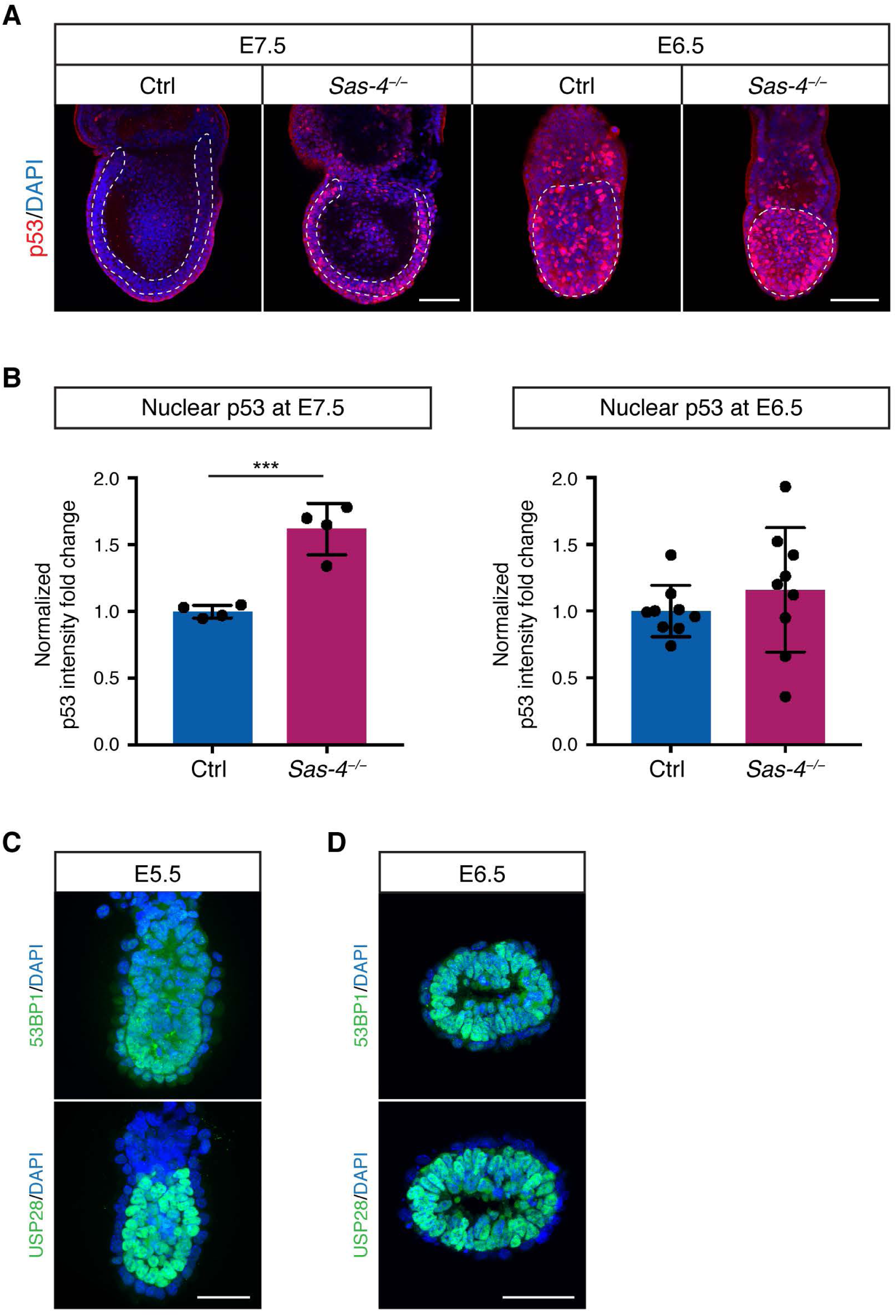
p53 upregulation in *Sas-4*^−/−^ embryos is evident by E7.5 and is not due to the onset of 53BP1 or USP28 expression. (A) Whole-mount immunostaining for p53 on control (Ctrl) and *Sas-4*^−/−^ embryos at E7.5 and E6.5. Representative sagittal planes are shown and the dotted lines demarcate the epiblast. Scale bars = 100 μm. (B) Quantification of nuclear p53 fluorescence intensity in the epiblast, normalized to control embryos in the same batch at E7.5 (n = 4) and at E6.5 (n = 9). *** p < 0.001. Error bars represent mean ± s.d. (C) Sagittal planes of whole-mount immunostaining for 53BP1 and USP28 on WT embryos at E5.5. Scale bar = 50 μm. (D) Immunostaining for 53BP1 and USP28 on sagittal sections of WT embryos at E6.5. Scale bar = 50 μm.

### USP28 and 53BP1 are expressed in the epiblast before E6

We next asked whether the upregulation of p53 in *Sas-4*^−/−^ embryos around E7, and not before, coincided with the onset of expression of either 53BP1 or USP28, the upstream regulators of p53. We performed immunostaining of 53BP1 and USP28 in control and *Sas-4*^−/−^ embryos at E5.5 and E6.5. Both 53BP1 and USP28 were expressed in WT embryos at E5.5 (Fig. 2C) and E6.5 (Fig. 2D). Of note, USP28 expression was clearly detectable in the embryonic epiblast but not in the surrounding visceral endoderm. *Sas-4*^−/−^ embryos also expressed both 53BP1 and USP28 at E6.5 (Fig. EV2C). The data indicated that the regulation of the onset of 53BP1 or USP28 expression does not seem to be responsible for p53 upregulation and activation of the mitotic surveillance pathway in *Sas-4*^−/−^ embryos, suggesting that other mechanisms establish the pathway around E7.

### The proper growth of *Sas-4*^−/−^ mESCs is dependent on p53

To study the dynamics of the mitotic surveillance pathway activation, we derived primary mESCs from WT and *Sas-4*^−/−^ blastocysts at E3.5. *Sas-4*^−/−^ primary mESCs were successfully derived and propagated *in vitro* and lacked detectable centrosomes in interphase cells, as judged by γ-tubulin (TUBG) staining, compared to WT cells, which had centrosomes in every cell (Fig. 3A). TUBG aggregates were seen only at the poles of mitotic cells in *Sas-4*^−/−^ mESCs, consistent with our findings in *Sas-4*^−/−^ embryos where these PCM aggregates lacked centrioles (Bazzi & Anderson, 2014a). Both WT and *Sas-4*^−/−^ primary mESCs showed high levels of nuclear NANOG in media containing LIF and 2i, indicating their pluripotent potential (Fig. 3B). In pluripotent conditions (LIF and 2i), WT and *Sas-4*^−/−^ primary mESCs had seemingly similar levels of p53, as judged by immunofluorescence (Fig. 3B). Because *Sas-4*^−/−^ embryos upregulated p53 starting after E6.5 (Fig. 2), we reasoned that *Sas-4*^−/−^ mESC partial differentiation may trigger a similar response *in vitro*. Thus, we removed the pluripotency factors (LIF and 2i) for three days, and the pluripotency potential declined as shown by the decrease in NANOG nuclear signal in both WT and *Sas-4*^−/−^ mESCs (Fig. 3B). The partially differentiated mESCs are not likely to represent a specific lineage because mESCs first move into a transitional state as they exit self-renewal (Martello & Smith, 2014). Importantly, upon partial differentiation, nuclear p53 levels decreased in WT but not in *Sas-4*^−/−^ mESCs (Fig. 3B). Quantification of the normalized nuclear p53 levels revealed that they were slightly, but significantly, higher in *Sas-4*^−/−^ mESCs compared to WT mESCs in pluripotent conditions, and this difference appeared more pronounced upon partial differentiation (Fig. 3C). Also, the decrease in p53 in WT mESCs upon partial differentiation was also significant (Fig. 3C).

**Fig. 3.**
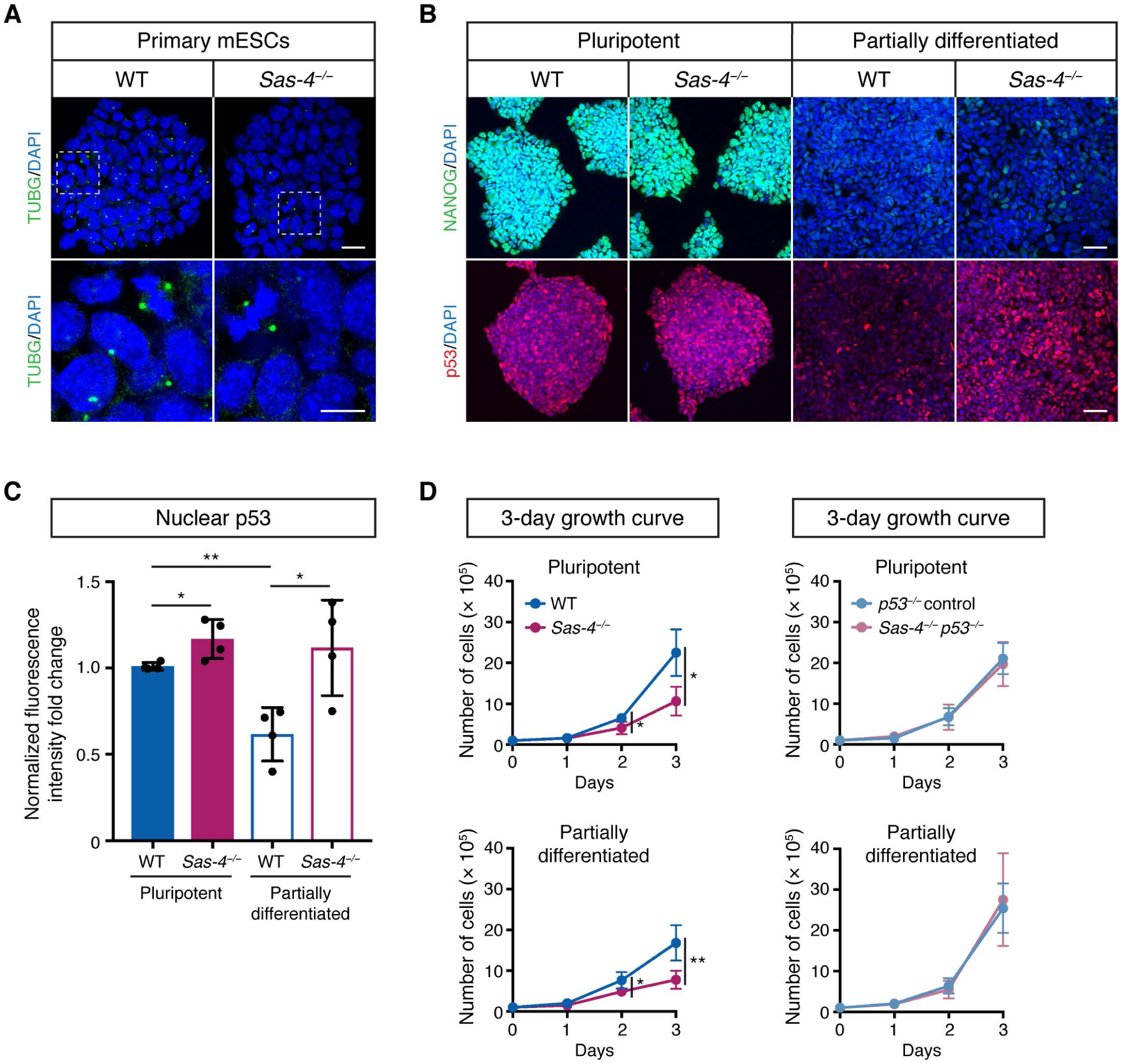
*Sas-4*^−/−^ mESCs activate the mitotic surveillance pathway. (A) Immunostaining for the centrosome marker γ-tubulin (TUBG) on WT and *Sas-4*^−/−^ primary mESCs. The bottom panels are magnifications of the areas marked in the top panels. Scale bars = 20 μm (top) and 10 μm (bottom). (B) Immunostaining for NANOG and p53 on WT and *Sas-4*^−/−^ primary mESCs in pluripotent and partially differentiated conditions. Scale bars = 50 μm. (C) Quantification of p53 fluorescence intensities shown in B normalized to WT (n = 4). ** p < 0.01, * p < 0.05. Error bars represent mean ± s.d. (D) Three-day growth curves of WT, *Sas-4*^−/−^, *p53*^−/−^, and *Sas-4*^−/−^ *p53*^−/−^ primary mESCs inthe indicated conditions starting with 10^5^ cells on Day 0 (n = 4 for each). ** p < 0.01, * p < 0.05. Error bars represent mean ± s.d.

Although *Sas-4*^−/−^ mESCs could be derived and propagated in pluripotent condition cultures, we noticed that they grew slower than WT mESCs (Fig. 3D). The growth defect became more obvious upon partial differentiation (Fig. 3D). To check whether the slower growth in *Sas-4*^−/−^ mESCs was dependent on p53 and the possible activation of the mitotic surveillance pathway, we generated *Sas-4*^−/−^ *p53*^−/−^ and *p53*^−/−^ control mESCs using CRISPR/Cas9 (see Methods and Fig. EV3). The data showed that *Sas-4*^−/−^ *p53*^−/−^ completely rescued the growth delay phenotype relative to *p53*^−/−^ and WT mESCs under pluripotent and partially differentiated conditions (Fig. 3D).

### Mitotic surveillance pathway activation is associated with prolonged mitosis *in vivo* and *in vitro*

We have previously shown that prometaphase was prolonged in *Sas-4*^−/−^ embryos at E7.5 and at E8.5 (Bazzi & Anderson, 2014a). To address whether the activation of the mitotic surveillance pathway around E7 coincided with the onset of prolonged mitosis in *Sas-4*^−/−^ embryos, we performed immunostaining for the mitotic marker phospho-histone H3 (pHH3) at E6.5. We calculated the mitotic index, the percentage of pHH3-positive cells in the epiblast, as an indirect measure of mitotic duration and detected no difference between control and *Sas-4*^−/−^ embryos at E6.5 (Fig. 4A). In contrast, our previous data showed that the mitotic index of *Sas-4*^−/−^ embryos at E7.5 was significantly higher than that of control embryos (Fig. 4A) (Bazzi & Anderson, 2014a). The data indicated that the mitotic surveillance pathway activation through p53 upregulation temporally correlates with prolonged mitosis *in vivo*.

**Fig. 4.**
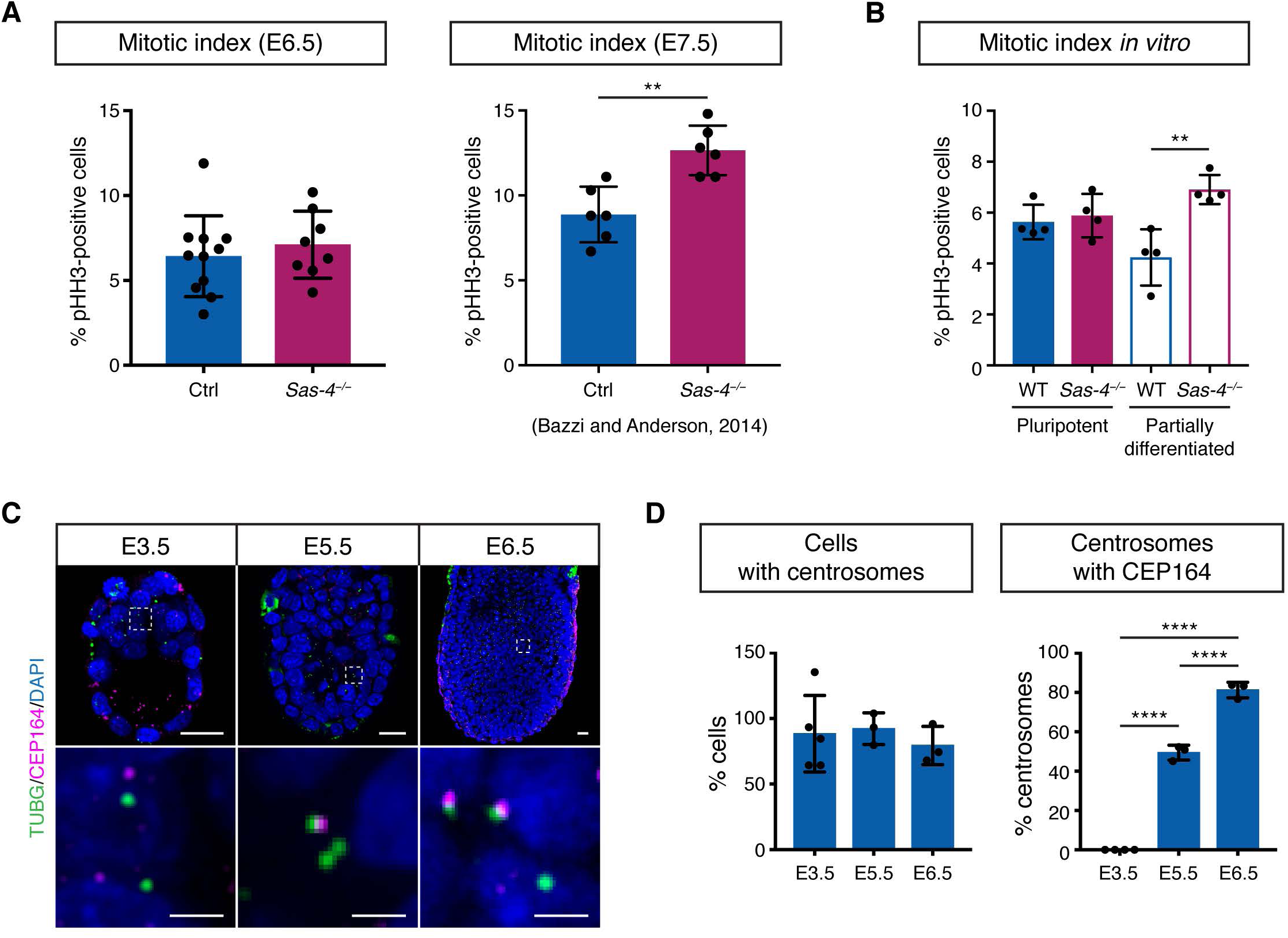
Changes in the mitotic index in developing *Sas-4*^−/−^ embryos correlate with centriole maturation. (A, B) The mitotic index, or percentage of pHH3-positive cells, of control and *Sas-4*^−/−^ embryos at E6.5 (n = 8) and E7.5 (n = 6) (B) and mESCs (n = 4) in different culture conditions (C). The graph at E7.5 represents our previously published data (Bazzi and Anderson, 2014). ** p < 0.01. Error bars represent mean ± s.d. (C) Sagittal planes of immunostaining for TUBG and CEP164 on WT mouse embryos from E3.5 to E6.5. The bottom panels are magnifications of the areas marked in the top panels. Scale bars = 20 μm (top) and 3 μm (bottom). (D) Quantification of the percentage of epiblast cells with centrosomes (TUBG) and the percentage of centrosomes with CEP164. **** p < 0.0001. Error bars represent mean ± s.d.

In line with the embryo data *in vivo*, the mitotic indices of WT and *Sas-4*^−/−^ mESCs *in vitro* were similar in pluripotency. However, upon partial differentiation, the mitotic index of *Sas-4*^−/−^ mESCs was significantly higher than that of WT mESCs (Fig. 4B). These findings indicated that the enhanced activation of the mitotic surveillance pathway in mESCs also correlates with prolonged mitosis upon partial differentiation, and that the growth dynamics of *Sas-4*^−/−^ mESCs largely resemble those of *Sas-4*^−/−^ embryos.

### Gradual centriole maturation correlates with the establishment of the mitotic surveillance pathway *in vivo*

We next hypothesized that the centrioles that are first formed by *de novo* biogenesis around E3 were not fully mature yet and that their maturation correlates with the delayed response to centriole loss in *Sas-4*^−/−^ embryos around E7. Therefore, we performed immunostaining for TUBG and the distal appendage protein CEP164, as a marker of the more mature mother centrioles, on developing embryos between E3.5 and E6.5. Starting at E3.5, almost all the cells contained centrosomes marked by TUBG foci (Fig. 4C, D). Intriguingly, CEP164 did not localize to these centrosomes at E3.5, supporting our hypothesis that the centrioles were not mature (Fig. 4C, D). At E5.5, around 65% of the centrosomes in the epiblast colocalized with CEP164, and the percentage increased to 85% at E6.5 (Fig. 4C, D). We concluded that the newly formed centrioles in mouse embryos gradually mature to participate in mitosis and cilia formation overlapping with the activation of the mitotic surveillance pathway in *Sas-4*^−/−^ centriole mutant embryos around E7 (Fig. 5).

**Fig. 5.**
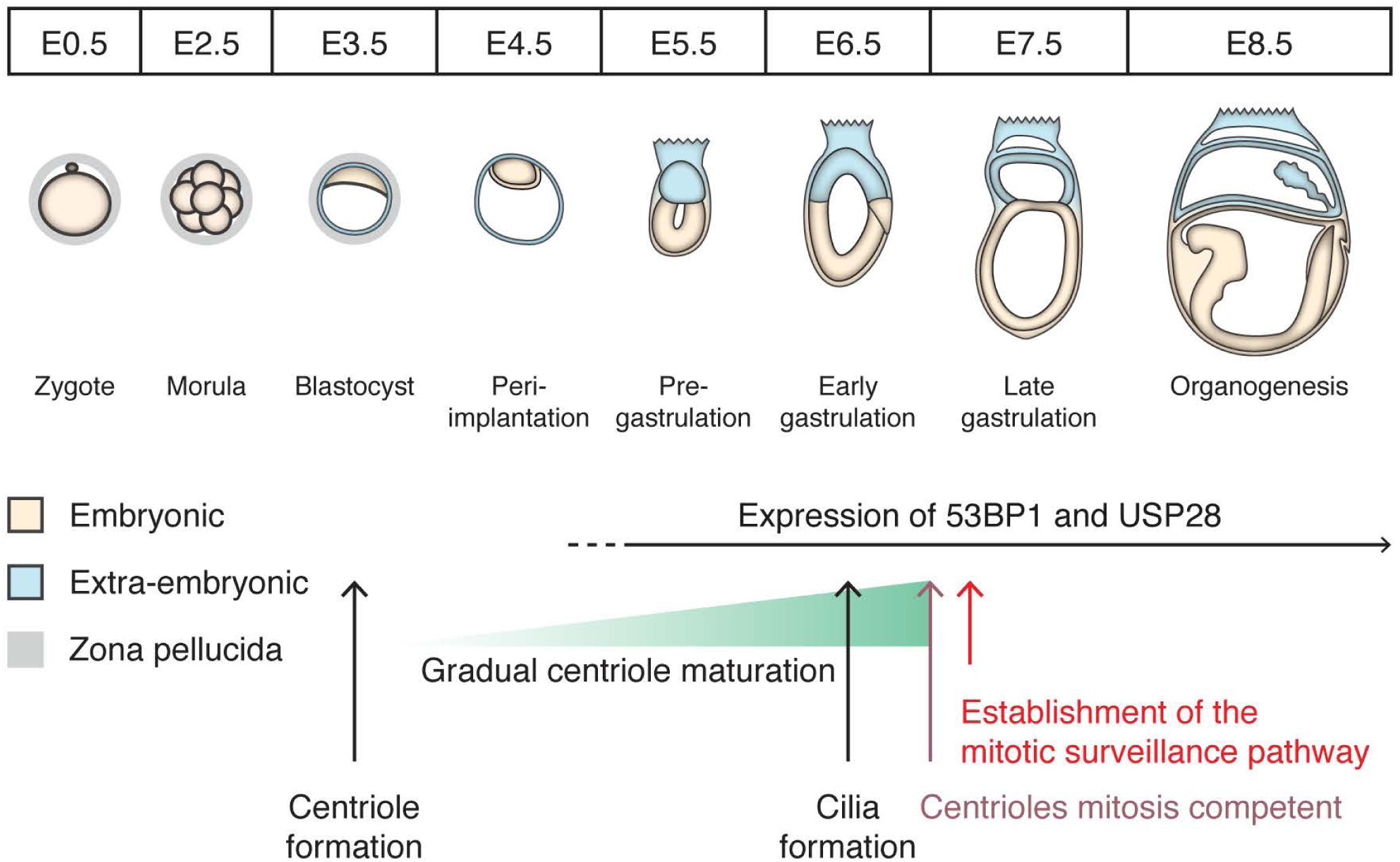
A schematic model of the correlation between gradual centriole maturation and centrosome functions during mouse embryonic development. Centrioles first form by *de novo* biogenesis around E3 and gradually mature to provide a template for cilia and MTOCs for mitosis around E7, when the mitotic surveillance pathway is established.

Although mitosis is usually the shortest phase of the cell cycle and lasts only around half an hour, it is an essential phase where the segregation of DNA and other cellular components must be precisely accomplished. In addition to the well-studied spindle assembly checkpoint (SAC), mammalian cells have developed a newly discovered pathway to monitor mitosis termed the mitotic surveillance pathway that is independent of the SAC (Lambrus & Holland, 2017). This pathway seems to be limited to mammalian systems because organisms such as *Drosophila melanogaster* lack 53PB1 and USP28 homologs (Lambrus & Holland, 2017), and zygotic *Sas-4* mutant flies survive until adulthood (Basto *et al*, 2006).

Both control and *Sas-4*^−/−^ embryos show relatively high nuclear p53 around E6.5, which has been reported in WT embryos at E5.5 and E6.5 (Bowling *et al*, 2018). It has been suggested that p53 may be involved in cellular competition during this stage of development to eliminate less fit cells before the germline is selected (Zhang *et al*, 2017). Higher p53 levels in *Sas-4*^−/−^ embryos around E7 coincide with the window of the initiation of gastrulation as well as the appearance of cilia on epiblast-derived lineages (Bangs *et al*., 2015). Our data largely exclude the lack of cilia *per se* (Fig. EV2) or the expression of the mitotic surveillance pathway components (Fig. 2C, D) as determinants of pathway activation after a lag period and at a specific developmental window. In line with this, 53BP1 expression has been reported throughout mouse pre-implantation development (Ziegler-Birling *et al*, 2009). In addition, USP28 expression was restricted to the epiblast, which may explain why the fast proliferating epiblast cells seem to be more affected by centriole loss compared to the visceral endoderm (Bazzi & Anderson, 2014a). Collectively, our data support a model whereby the newly formed centrioles around E3 gradually mature during development until around E7, when they are competent to participate in cilia formation as well as act as efficient MTOCs during mitosis (Fig. 5). As such, *Sas-4*^−/−^ centriole mutant embryos may not activate the p53-dependent mitotic surveillance pathway until centrioles are more mature and required for mitosis. This gradual transition is reminiscent of the earlier transition from meiotic- to mitotic-like divisions during pre-implantation and may be a general phenomenon in development including, for example, cilia formation, elongation and function (Bangs *et al*., 2015; Courtois & Hiiragi, 2012).

## Materials and Methods

### Animals and genotyping

The *Sas-4*^−/−^ mice (*Cenpj*^*tm1d(EUCOMM)Wtsi/tm1d(EUCOMM)Wtsi*^) (Bazzi & Anderson, 2014a) and the *Ift88*^−/−^ null mouse allele generated from the *Ift88*^*fl/fl*^ allele (*Ift88*^*tm1Bky*^) (Haycraft *et al*., 2007) were used in this study. The CRISPR/Cas9 endonuclease-mediated knockouts of *53bp1*^−/−^ and *Usp28*^−/−^ were generated by the CECAD *in vivo* Research Facility using microinjection or electroporation of the corresponding gRNA, Cas9 mRNA and Cas9 protein into fertilized zygotes (Table 1) (Chu *et al*, 2016; Troder *et al*, 2018). The gRNA target sequence predictor tool developed by the Broad Institute was used to design gRNAs (Doench *et al*, 2016).

**Table 1.**
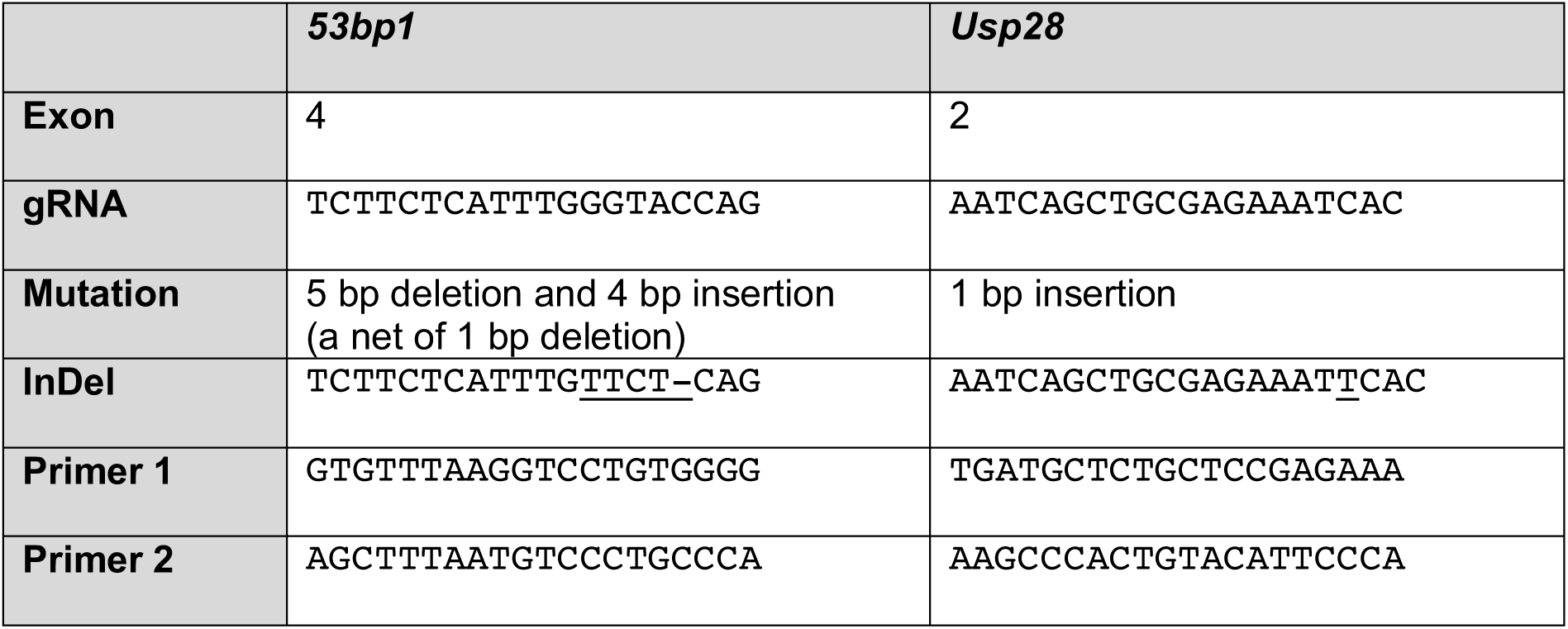
Information for CRISPR/Cas9 and genotyping of *53bp1* and *Usp28* mouse alleles.

The animals were housed and bred under standard conditions in the CECAD animal facility, and the allele generation (84-02.04.2014.A372) and experiments (84-02.05.50.15.039) were approved by the Landesamt für Natur, Umwelt, und Verbraucherschutz Nordrhein-Westfalen (LANUV-NRW) in Germany. All the phenotypes were analyzed in the FVB/NRj background. Genotyping was carried out using standard and published PCR protocols. The PCR products for *53bp1*- and *Usp28-*mutant mice were digested with *Kpn*I and *Apo*I restriction enzymes (New England BioLabs; Ipswich, MA, USA), respectively, to distinguish the WT and mutant alleles.

### Mouse embryonic stem cell culture

Primary mESCs were derived from E3.5 blastocysts as previously described (Bryja *et al*, 2006), cultured on feeder cells that were proliferation-inactivated with mitomycin C (Sigma Aldrich; St. Louis, MO, USA) and 0.1% gelatin-coated plates (Sigma Aldrich). They were maintained in media containing Knock-Out DMEM (Thermo Fisher Scientific; Waltham, MA, USA), supplemented with 15% Hyclone fetal bovine serum (FBS; VWR; Radnor, PA, USA), 2 mM L-glutamine (Biochrom; Berlin, Germany), 1% penicillin/streptomycin (Biochrom), 0.1 mM MEM non-essential amino acids (Thermo Fisher Scientific), 1 mM sodium pyruvate (Thermo Fisher Scientific), 0.1 mM β-mercaptoethanol (Thermo Fisher Scientific), 1000 U/ml leukemia inhibitory factor (LIF; Merck; Darmstadt, Germany), and with 1 μM PD0325901 (Miltenyi Biotec; Bergisch Gladbach, Germany) and 3 μM CHIR99021 (Miltenyi Biotec), together abbreviated as 2i. Primary mESCs were gradually weaned off feeder cells and maintained in feeder-free conditions. To induce partial differentiation, feeder-free primary mESCs were split and cultured in media without LIF and 2i for three days.

### Generating CRISPR-modified primary mESCs

The gRNA sequence for targeting *p53* was cloned as double-stranded DNA oligonucleotides into the *Bbs*I restriction site of the pX330-U6-Chimeric_BB-CBh-hSpCas9 vector (Addgene; Watertown, MA, USA) modified with a Puro-T2K-GFP cassette containing puromycin-resistance and eGFP expression by Dr. Leo Kurian’s research group (Center for Molecular Medicine Cologne).

*p53*^−/−^ and *Sas-4*^−/−^ *p53*^−/−^ mESCs (Table 2) were generated by lipofection of the modified pX330 vector containing the gRNA target sequences using Lipofectamine 3000 (Thermo Fisher Scientific). One day after transfection, 2 μg/ml puromycin (Sigma Aldrich) was added to the medium for two days, and the cells were allowed to recover in regular medium up to one week after transfection. Single colonies were picked under a dissecting microscope and were expanded. *p53* null cell lines were confirmed with sequencing (primers in Table 2), immunofluorescence, and western blotting.

**Table 2.**
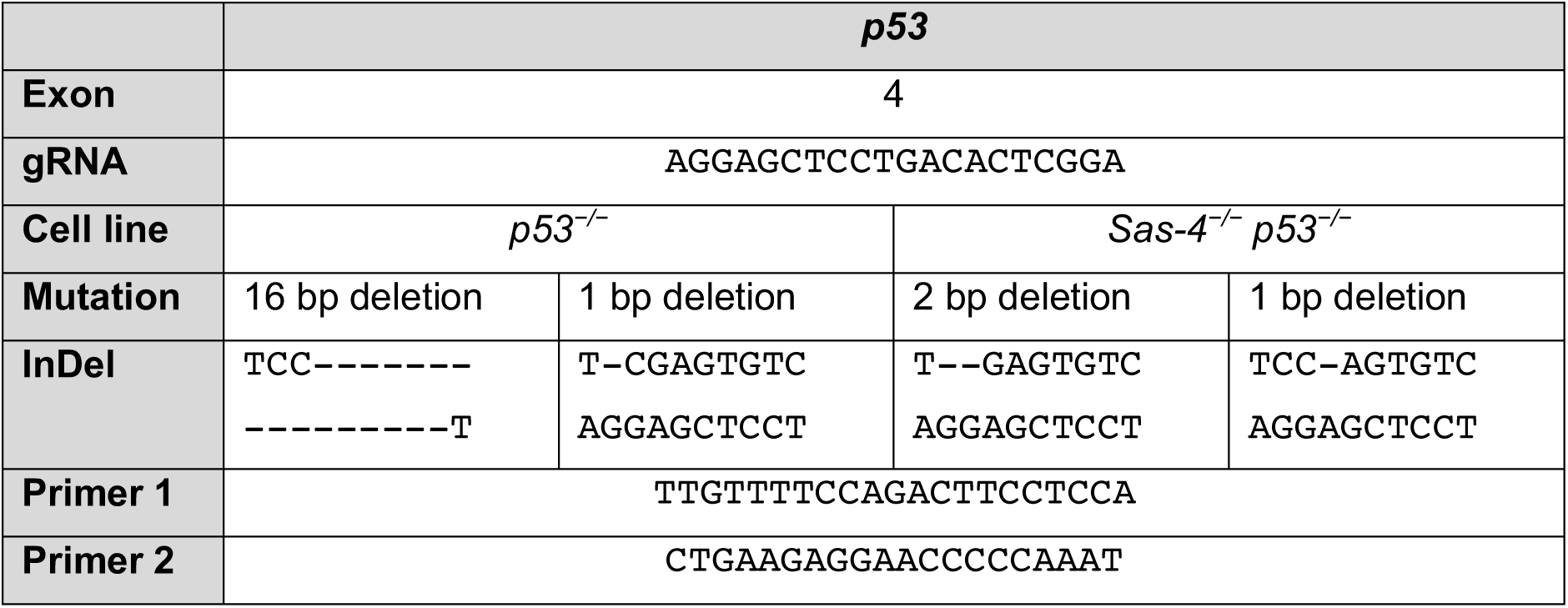
Information for CRISPR/Cas9-generated *p53* alleles in mESCs.

### Growth assay

To determine the growth kinetics of mESCs over three days, WT, *Sas-4*^−/−^, *p53*^−/−^, and *Sas-4*^−/−^ *p53*^−/−^ mESCs were seeded at 10^5^ cells per well of a 6-well plate in media with or without LIF and 2i in triplicate. One set was counted every day for three days using a hemocytometer. Two pairs of each genotype from separate derivations were counted twice and constituted four biological replicates.

### Embryo dissection, immunofluorescence and imaging

Pregnant female mice (E3.5 and E9.5) were sacrificed by cervical dislocation for embryo dissections under a dissecting microscope (M165C or M80, Leica Microsystems; Wetzlar, Germany) as previously described (Behringer *et al*, 2014; Bryja *et al*., 2006). The embryos were fixed in 4% paraformaldehyde (PFA; Carl Roth; Karlsruhe, Germany) for 2 h at room temperature or overnight at 4°C. Embryos at E3.5 were fixed in 4% PFA for 30 min and in ice cold methanol for 15 min. After fixing, the embryos were washed with phosphate buffered saline (PBS; VWR), and then either used for whole-mount immunostaining or cryoprotected in 30% sucrose overnight at 4°C. The embryos were transferred from 30% sucrose and embedded in optimum cutting temperature (OCT) compound (Sakura Finetek; Alphen an den Rijn, Netherlands) for cryosectioning.

Whole-mount immunofluorescence staining of intact mouse embryos was performed as previously described (Xiao *et al*, 2018). The embryos were then mounted in 1% low-melting agarose (Lonza; Basel, Switzerland) on a glass bottom dish (Thermo Fisher Scientific), covered in VectaShield mounting medium (Linaris; Dossenheim, Germany), and kept cold and protected from light until imaging. After imaging, the embryos were removed from the agarose, washed and digested for genotyping. Embryos at E3.5 were directly imaged in PBS in a glass bottom dish.

OCT-embedded embryos were cryosectioned at 8 μm thickness and the slides were fixed with ice-cold methanol for 10 min at −20°C, then washed two times with wash buffer containing 0.2% Triton-X in PBS while shaking and blocked with wash buffer with 5% heat-inactivated goat serum for 1 h at room temperature. The slides were incubated with primary antibodies diluted in blocking solution overnight at 4°C. After washing two times with wash buffer, the slides were incubated with secondary antibodies and DAPI in blocking solution for 1 h at room temperature. After washing, the slides were mounted with coverslips using Prolong Gold (Cell Signaling Technology; Danvers, MA, USA).

For immunofluorescence of mESCs, 2×10^4^ cells were seeded per chamber onto pre-gelatinized Lab-Tek II chamber slides (Thermo Fisher Scientific). After three days of culturing in corresponding media, the cells were washed with PBS, fixed with 4% PFA for 10 min at room temperature and washed three times with PBS. Next, the cells were fixed with ice-cold methanol for 10 min at −20°C, permeabilized with 0.5% Triton-X in PBS for 5 min at room temperature, and blocked with 5% heat-inactivated goat serum (Thermo Fisher Scientific) for at least 15 min at room temperature. The cells were incubated with primary antibodies diluted in blocking solution overnight at 4°C. After washing three times with wash buffer, the cells were incubated with secondary antibodies and DAPI diluted 1:1000 in blocking solution for 1 h at room temperature. After washing, the chamber was removed from the glass slide and coverslips were mounted using Prolong Gold anti-fade reagent (Cell Signaling Technology). The images were obtained using an SP8 confocal microscope (Leica Microsystems).

### Antibodies

Primary antibodies used in this study and their dilutions and sources are listed in Table 3. The secondary antibodies used were Alexafluor® 488, 568, or 647 conjugates (Life Technologies) and diluted at 1:1000, in combination with DAPI (AppliChem) at 1:1000.

**Table 3.**
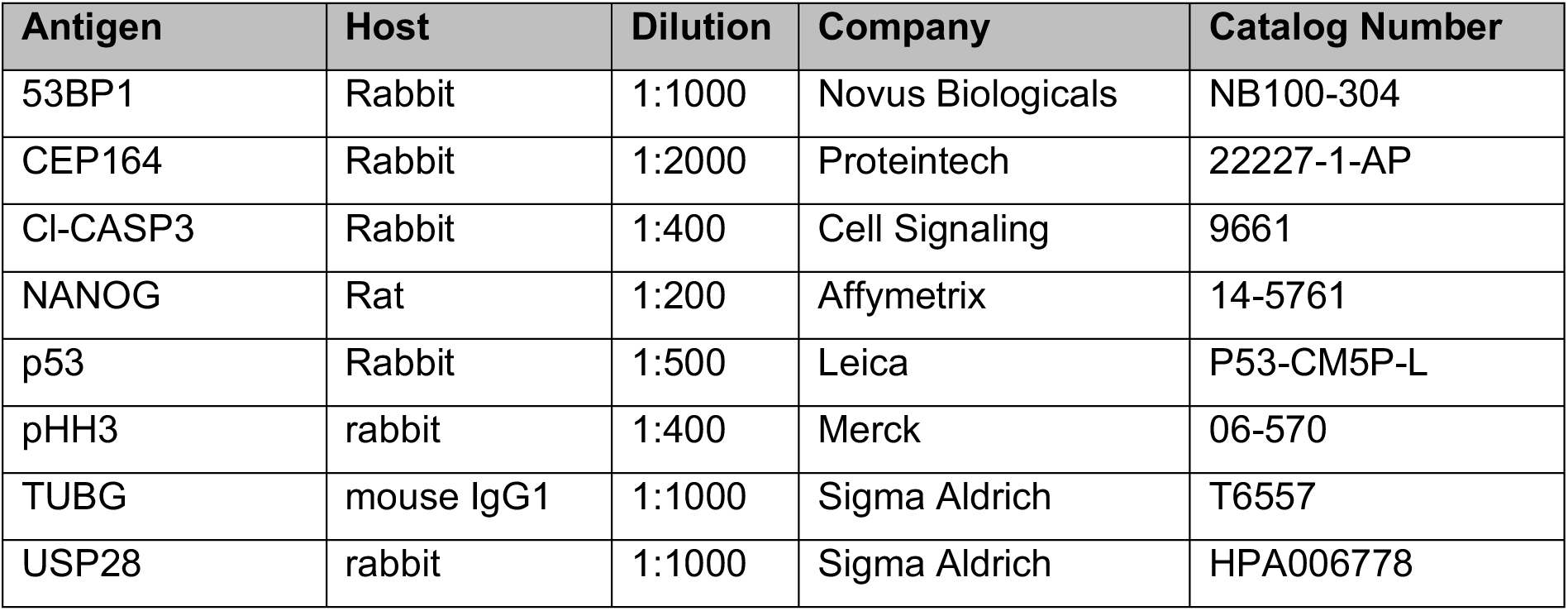
List of primary antibodies used in this study.

### Image analyses

Images of whole E6.5 or E7.5 embryos or mESCs stained with p53 or pHH3 and DAPI were quantified using ImageJ (NIH, Maryland, USA). A maximum projection image of mESCs or the middle five slices of the embryo were generated. The DAPI channel was used to set a threshold to obtain a region of interest. p53 and DAPI fluorescence intensities were measured, and the p53 intensity was normalized to the DAPI intensity. The average p53 intensity of controls was set to 1.0 and a fold-change p53 intensity was calculated. Centrosomes (defined as one TUBG focus or two close TUBG foci) and CEP164 foci were manually quantified using ImageJ (NIH). The number of nuclei were quantified using the image-based tool for counting nuclei (ITCN) ImageJ plug-in.

### Western Blotting

mESCs were scraped in radioimmunoprecipitation assay (RIPA) buffer containing 150 mM NaCl, 50 mM Tris pH 7.6, 1% Triton X-100 (Sigma-Aldrich), 0.25% sodium deoxycholate, and 0.1% sodium dodecyl sulfate (SDS; AppliChem; Darmstadt, Germany) with an ethylenediaminetetraacetic acid (EDTA)-free protease inhibitor cocktail (Merck), phosphatase inhibitor cocktail sets II (Merck) and IV (Merck), and phenylmethylsulfonyl fluoride (PMSF; Sigma-Aldrich). Protein concentration was measured with an RC DC protein assay kit (Bio-Rad; Feldkirchen, Germany). 10 μg protein per sample was loaded. SDS-polyacrylamide gel electrophoresis (PAGE) and immunoblotting were performed following standard procedures (Kurien and Scofield, 2006; Towbin et al., 1979). Following SDS-PAGE, the proteins were transferred to polyvinylidene fluoride (PVDF) membranes (Merck) that were activated in methanol (Carl Roth) for 1 min, blocked in 5% milk (Carl Roth) for 1 h, and incubated with an anti-p53 antibody (1:5000; Leica Biosystems; Buffalo Grove, IL, USA) or an anti-GAPDH antibody (1:10^4^; Merck) overnight at 4°C. The membranes were washed with Tris buffered saline containing Tween 20 (AppliChem; TBST) and incubated with secondary anti-rabbit (GE Healthcare; Chicago, IL, US) or anti-mouse (GE Healthcare) antibodies linked with horseradish peroxidase (HRP) at 1:10^4^ for 1 h at room temperature. Finally, the membranes were washed with TBST and incubated with enhanced chemiluminescence (ECL; GE Healthcare; Chicago, IL, USA) and signals were detected with on film (GE Healthcare) in a dark room.

### Statistical Analyses

Statistical analyses comparing two groups of data using a two-tailed Student’s t-test with a cutoff for significance of less than 0.05 and graphic representations with standard deviations were performed using GraphPad Prism 7 (GraphPad Software, San Diego, CA, USA).

## Acknowledgements

We thank the CECAD *in vivo* research facility for generating and maintaining our mouse lines and the CECAD imaging facility for microscopy support. The work was supported by the Deutsche Forschungsgemeinschaft (DFG, German Research Foundation) [BA 5810/1-1 to H.B]. The funders had no role in study design, data collection and analysis, decision to publish, or preparation of the manuscript.

## Author contributions

Conceptualization: C.X. and H.B.; Methodology: C.X., M.G., C.G., M.M., R.F. and H.B.; Software: C.X., M.G., C.G.; Formal Analysis: C.X, M.G., C.G.; Investigation: C.X., M.G., C.G., M.M., R.F. and H.B.; Writing: C.X. and H.B.; Visualization: C.X., M.G., C.G.; Supervision, Project administration and Funding Acquisition: H.B.

## Expanded View Figure Legends

**Fig. EV1.**
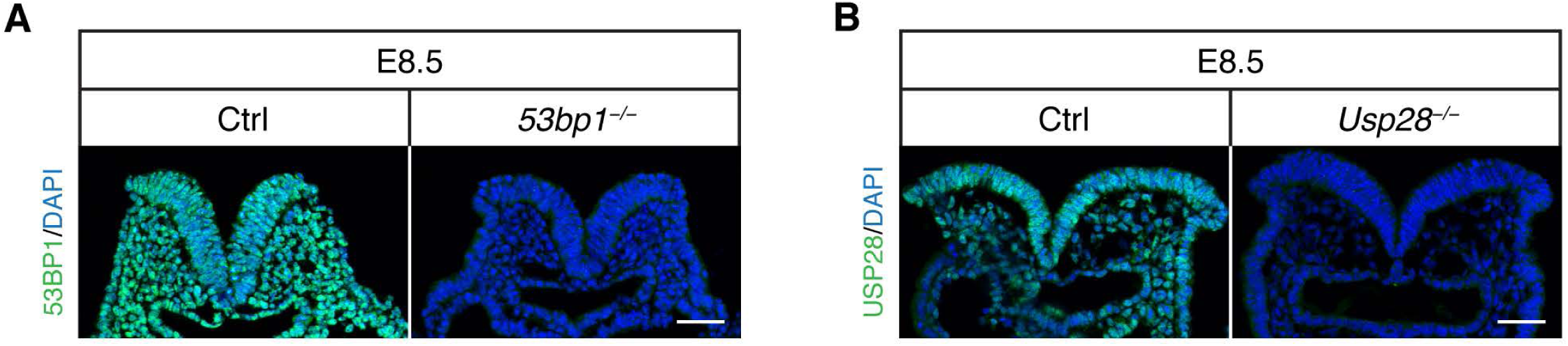
*53bp1*^−/−^ and *Usp28*^−/−^ are null alleles. (A, B) Immunostaining for 53BP1 (A) or USP28 (B) on sagittal sections of control (Ctrl) and *Sas-4*^−/−^ *53bp1*^−/−^ (A) or *Usp28*^−/−^ (B) embryos at E8.5. The signals for the corresponding proteins are not detectable in the mutants compared to controls. The V-shaped neural plate, underlying mesenchyme and gut tube are shown. Scale bars = 50 μm.

**Fig. EV2.**
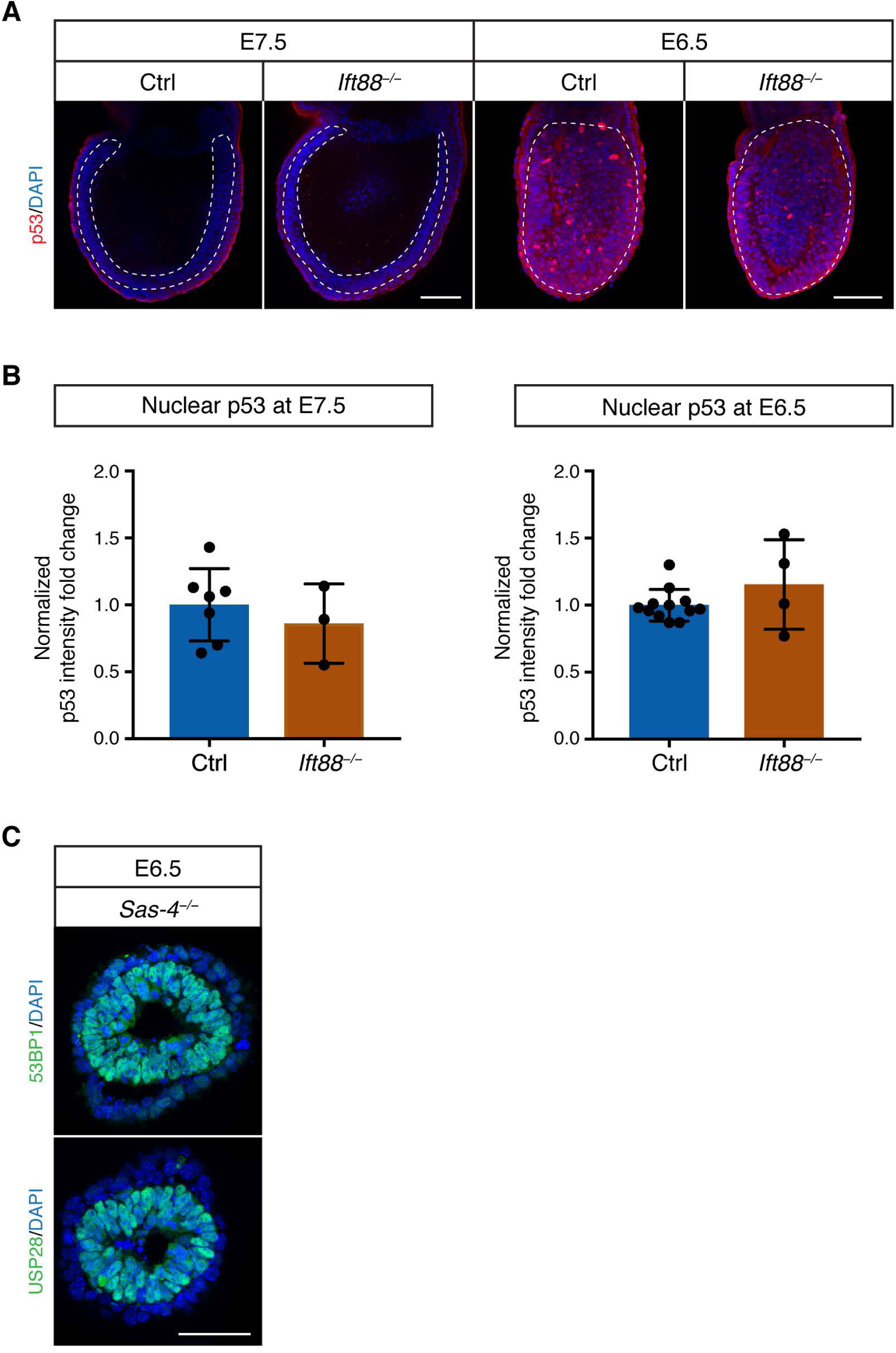
*Ift88*^−/−^ cilia mutants do not upregulate p53, and both 53BP1 and USP28 are expressed in *Sas-4*^−/−^ embryos at E6.5. (A) Immunostaining for p53 on whole-mount Ctrl and *Ift88*^−/−^ embryos at E7.5 and E6.5. Mid-sagittal planes are shown with the dotted lines demarcating the epiblast. Scale bars = 100 μm. (B) Quantification of p53 nuclear fluorescence intensity in the epiblast normalized to Ctrl embryos in the same batch at E7.5 (n = 3) and E6.5 (n = 4). Error bars represent mean ± s.d. (C) Immunostaining for 53BP1 and USP28 on transverse sections of *Sas-4*^−/−^ embryos at E6.5. Scale bar = 50 μm.

**Fig. EV3.**
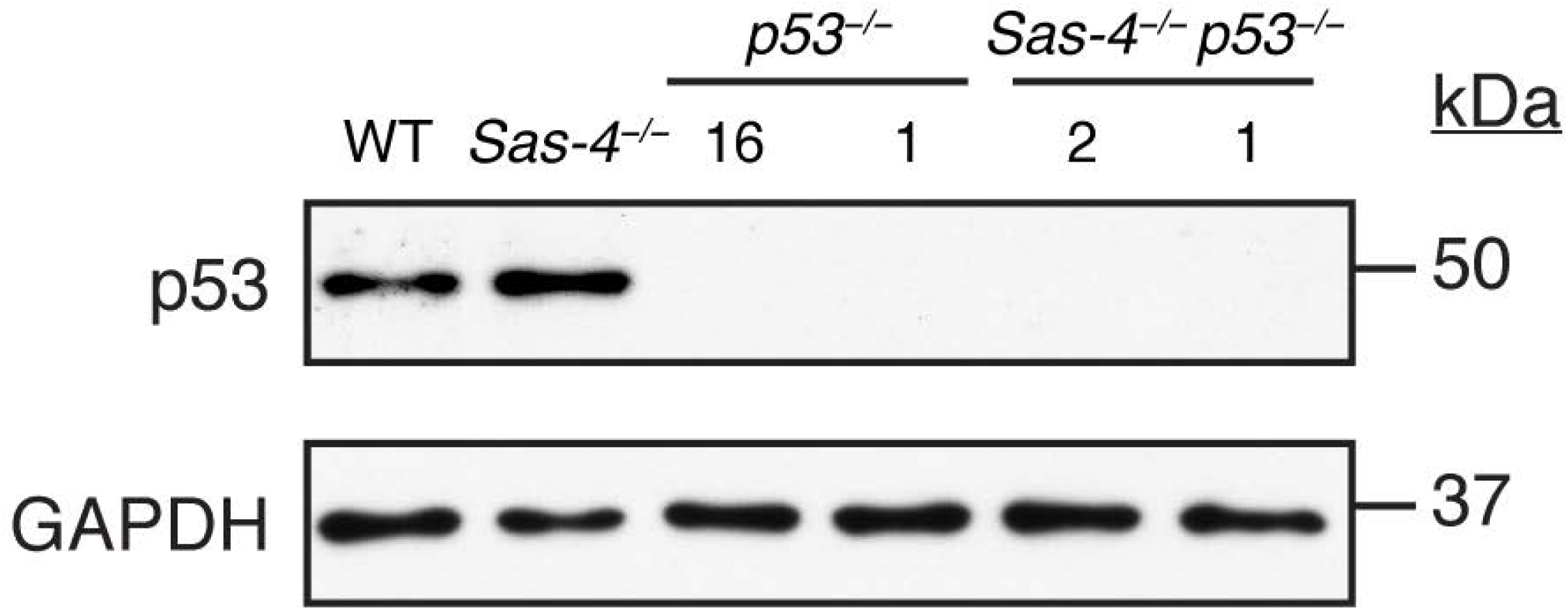
*p53*^−/−^ and *Sas-4*^−/−^ *p53*^−/−^ primary mESCs are null alleles for p53. Western blot analysis for p53 and GAPDH loading control on WT, *Sas-4*^−/−^, *p53*^−/−^, and *Sas-4*^−/−^ *p53*^−/−^ mESC lysates. The numbers below *p53*^−/−^ and *Sas-4*^−/−^ *p53*^−/−^ indicate the number of base pairs deleted.

## References

Bangs FK, Schrode N, Hadjantonakis AK, Anderson KV (2015) Lineage specificity of primary cilia in the mouse embryo. Nat Cell Biol 17: 113–122

Basto R, Lau J, Vinogradova T, Gardiol A, Woods CG, Khodjakov A, Raff JW (2006) Flies without centrioles. Cell 125: 1375–1386

Bazzi H, Anderson KV (2014a) Acentriolar mitosis activates a p53-dependent apoptosis pathway in the mouse embryo. Proc Natl Acad Sci U S A 111: E1491–1500

Bazzi H, Anderson KV (2014b) Centrioles in the mouse:cilia and beyond. Cell Cycle 13: 2809

Behringer R, Gertsenstein M, Vintersen Nagy K, Nagy A, 2014. Manipulating the Mouse Embryo: A Laboratory Manual, Fourth Edition, Cold Harbor Laboratory Press. pp. 198-200,268–271.

Bornens M (2012) The centrosome in cells and organisms. Science 335: 422–426

Bowling S, Di Gregorio A, Sancho M, Pozzi S, Aarts M, Signore M, M Ds, Martinez-Barbera JP, Gil J, Rodriguez TA (2018) P53 and mTOR signalling determine fitness selection through cell competition during early mouse embryonic development. Nat Commun 9: 1763

Bryja V, Bonilla S, Arenas E (2006) Derivation of mouse embryonic stem cells. Nat Protoc 1: 2082–2087

Chu VT, Weber T, Graf R, Sommermann T, Petsch K, Sack U, Volchkov P, Rajewsky K, Kuhn R (2016) Efficient generation of Rosa26 knock-in mice using CRISPR/Cas9 in C57BL/6 zygotes. BMC Biotechnol 16: 4

Conduit PT, Wainman A, Raff JW (2015) Centrosome function and assembly in animal cells. Nat Rev Mol Cell Biol 16: 611–624

Courtois A, Hiiragi T (2012) Gradual meiosis-to-mitosis transition in the early mouse embryo. Results and problems in cell differentiation 55: 107–114

Courtois A, Schuh M, Ellenberg J, Hiiragi T (2012) The transition from meiotic to mitotic spindle assembly is gradual during early mammalian development. J Cell Biol 198: 357–370

Doench JG, Fusi N, Sullender M, Hegde M, Vaimberg EW, Donovan KF, Smith I, Tothova Z, Wilen C, Orchard R et al (2016) Optimized sgRNA design to maximize activity and minimize off-target effects of CRISPR-Cas9. Nat Biotechnol 34: 184–191

Fong CS, Mazo G, Das T, Goodman J, Kim M, O’Rourke BP, Izquierdo D, Tsou MF (2016) 53BP1 and USP28 mediate p53-dependent cell cycle arrest in response to centrosome loss and prolonged mitosis. Elife 5

Graser S, Stierhof YD, Lavoie SB, Gassner OS, Lamla S, Le Clech M, Nigg EA (2007) Cep164, a novel centriole appendage protein required for primary cilium formation. J Cell Biol 179: 321–330

Gueth-Hallonet C, Antony C, Aghion J, Santa-Maria A, Lajoie-Mazenc I, Wright M, Maro B (1993) gamma-Tubulin is present in acentriolar MTOCs during early mouse development. Journal of Cell Science 105 (Pt 1): 157–166

Haycraft CJ, Zhang Q, Song B, Jackson WS, Detloff PJ, Serra R, Yoder BK (2007) Intraflagellar transport is essential for endochondral bone formation. Development 134: 307–316

Howe K, FitzHarris G (2013) A non-canonical mode of microtubule organization operates throughout pre-implantation development in mouse. Cell Cycle 12: 1616–1624

Kirkham M, Müller-Reichert T, Oegema K, Grill S, Hyman AA (2003) SAS-4 is a C. elegans centriolar protein that controls centrosome size. Cell 112: 575–587

Kleylein-Sohn J, Westendorf J, Le Clech M, Habedanck R, Stierhof YD, Nigg EA (2007) Plk4-induced centriole biogenesis in human cells. Dev Cell 13: 190–202

Kong D, Farmer V, Shukla A, James J, Gruskin R, Kiriyama S, Loncarek J (2014) Centriole maturation requires regulated Plk1 activity during two consecutive cell cycles. J Cell Biol 206: 855–865

Lambrus BG, Daggubati V, Uetake Y, Scott PM, Clutario KM, Sluder G, Holland AJ (2016) A USP28-53BP1-p53-p21 signaling axis arrests growth after centrosome loss or prolonged mitosis. J Cell Biol 214: 143–153

Lambrus BG, Holland AJ (2017) A New Mode of Mitotic Surveillance. Trends Cell Biol 27: 314–321

Lambrus BG, Uetake Y, Clutario KM, Daggubati V, Snyder M, Sluder G, Holland AJ (2015) p53 protects against genome instability following centriole duplication failure. J Cell Biol 210: 63–77

Leidel S, Gonczy P (2003) SAS-4 is essential for centrosome duplication in C elegans and is recruited to daughter centrioles once per cell cycle. Dev Cell 4: 431–439

Loncarek J, Khodjakov A (2009) Ab ovo or de novo? Mechanisms of centriole duplication. Mol Cells 27: 135–142

Manandhar G, Sutovsky P, Joshi HC, Stearns T, Schatten G (1998) Centrosome reduction during mouse spermiogenesis. Dev Biol 203: 424–434

Martello G, Smith A (2014) The nature of embryonic stem cells. Annu Rev Cell Dev Biol 30: 647–675

Meitinger F, Anzola JV, Kaulich M, Richardson A, Stender JD, Benner C, Glass CK, Dowdy SF, Desai A, Shiau AK et al (2016) 53BP1 and USP28 mediate p53 activation and G1 arrest after centrosome loss or extended mitotic duration. J Cell Biol 214: 155–166

Nichols J, Smith A (2011) The origin and identity of embryonic stem cells. Development 138: 3–8

Rosner MH, Vigano MA, Ozato K, Timmons PM, Poirier F, Rigby PW, Staudt LM (1990) A POU-domain transcription factor in early stem cells and germ cells of the mammalian embryo. Nature 345: 686–692

Siller SS, Sharma H, Li S, Yang J, Zhang Y, Holtzman MJ, Winuthayanon W, Colognato H, Holdener BC, Li FQ et al (2017) Conditional knockout mice for the distal appendage protein CEP164 reveal its essential roles in airway multiciliated cell differentiation. PLoS genetics 13: e1007128

Tang CJ, Fu RH, Wu KS, Hsu WB, Tang TK (2009) CPAP is a cell-cycle regulated protein that controls centriole length. Nat Cell Biol 11: 825–831

Troder SE, Ebert LK, Butt L, Assenmacher S, Schermer B, Zevnik B (2018) An optimized electroporation approach for efficient CRISPR/Cas9 genome editing in murine zygotes. PLoS One 13: e0196891

Williams RL, Hilton DJ, Pease S, Willson TA, Stewart CL, Gearing DP, Wagner EF, Metcalf D, Nicola NA, Gough NM (1988) Myeloid leukaemia inhibitory factor maintains the developmental potential of embryonic stem cells. Nature 336: 684–687

Wong YL, Anzola JV, Davis RL, Yoon M, Motamedi A, Kroll A, Seo CP, Hsia JE, Kim SK, Mitchell JW et al (2015) Cell biology. Reversible centriole depletion with an inhibitor of Polo-like kinase 4. Science 348: 1155–1160

Woolley DM, Fawcett DW (1973) The degeneration and disappearance of the centrioles during the development of the rat spermatozoon. Anat Rec 177: 289–301

Xiao C, Nitsche F, Bazzi H (2018) Visualizing the Node and Notochordal Plate In Gastrulating Mouse Embryos Using Scanning Electron Microscopy and Whole Mount Immunofluorescence. J Vis Exp

Ying QL, Wray J, Nichols J, Batlle-Morera L, Doble B, Woodgett J, Cohen P, Smith A (2008) The ground state of embryonic stem cell self-renewal. Nature 453: 519–523

Zhang G, Xie Y, Zhou Y, Xiang C, Chen L, Zhang C, Hou X, Chen J, Zong H, Liu G (2017) p53 pathway is involved in cell competition during mouse embryogenesis. Proc Natl Acad Sci U S A 114: 498–503

Ziegler-Birling C, Helmrich A, Tora L, Torres-Padilla ME (2009) Distribution of p53 binding protein 1 (53BP1) and phosphorylated H2A.X during mouse preimplantation development in the absence of DNA damage. Int J Dev Biol 53: 1003–1011

